# Two step PCR method to exchange the resistance cassette of a vector

**DOI:** 10.1101/2021.01.28.428618

**Authors:** Nils Cremer, Anne Diehl

**Affiliations:** Department of NMR-supported structural Biology, Leibniz-Forschungsinstitut für Molekulare Pharmakologie (FMP), Robert-Rössle-Str. 10, 13125 Berlin, Germany

**Keywords:** ligation independent cloning, exchange antibiotic resistance, protein expression

## Abstract

For co-transformation of two plasmids, both have to possess different antibiotic selection markers. If that is not the case, normally the gene of interest (GOI) is subcloned into another vector. Here we introduce a fast and easy method to exchange the antibiotic resistance cassette (ARC) in only two PCR steps.

**Method Summary:** To shuttle the antibiotic resistance cassette (ARC) from one vector to another, one can amplify the ARC of interest and use the resulting PCR-product as a primer pair for the next amplification step. Simply remove parental DNA template by *Dpn*I digestion, transform PCR product directly in *E. coli cells*, select transformants on an appropriate agar plate and isolate target vector by plasmid preparation.

---

Progress in structural biology depends on properly folded und functional recombinant proteins. After completion of the big structural genomics projects the low hanging fruits for *E. coli* expression have been almost entirely achieved. The challenge now is to produce difficult-to-express proteins in an improved *E. coli* environment, as this host is the easiest and cheapest one to handle with respect to NMR-labelling. One possibility is to optimize *E. coli* protein homeostasis by initiation of stress by sigma factor 32 (1). The appropriate plasmid pBAD.Sigma32.I54N (Figure 1A) carrying ampicillin resistance can be obtained via Addgene from Jeff Kelly. Sigma 32 causes the up-regulation of proteostasis network components like chaperones, folding enzymes or proteases. We have used that system surprisingly well for stable expression of an intrinsic unfolded protein with improved yield. Encouraged by that, we now want to use it for soluble expression of proteins with a high tendency to form inclusion bodies. Unfortunately most of our constructs have ampicillin as the selection marker, making co-transformation with pBAD.Sigma32.I54N impossible due to resistance being the same. Instead of re-cloning all our constructs we decided to change the antibiotic resistance cassette (ARC) of pBAD.Sigma32.I54N. A common pET26b vector (69862, Merck, Germany) was chosen as a template for the kanamycin ARC (Figure 1B). Initially both ARCs with their flanking regions were analyzed using alignments with respect to homolog regions in both directions. Indeed a reverse complimentary sequence downstream the ampicillin ARC of pBAD.Sigma32.I54N is identical to a sequence upstream the kanamycin ARC of pET26b and vice versa. Therefore two XL primers were synthesized (BioTez, Berlin, Germany) according to Table 1. With these primers the ampicillin ARC downstream of the sigma 32 and with the same translational orientation will be replaced by the kanamycin ARC in the opposite direction (Figure 1C). The reverse orientation is recommended for the kanamycin ARC (2).

**Figure 1.**
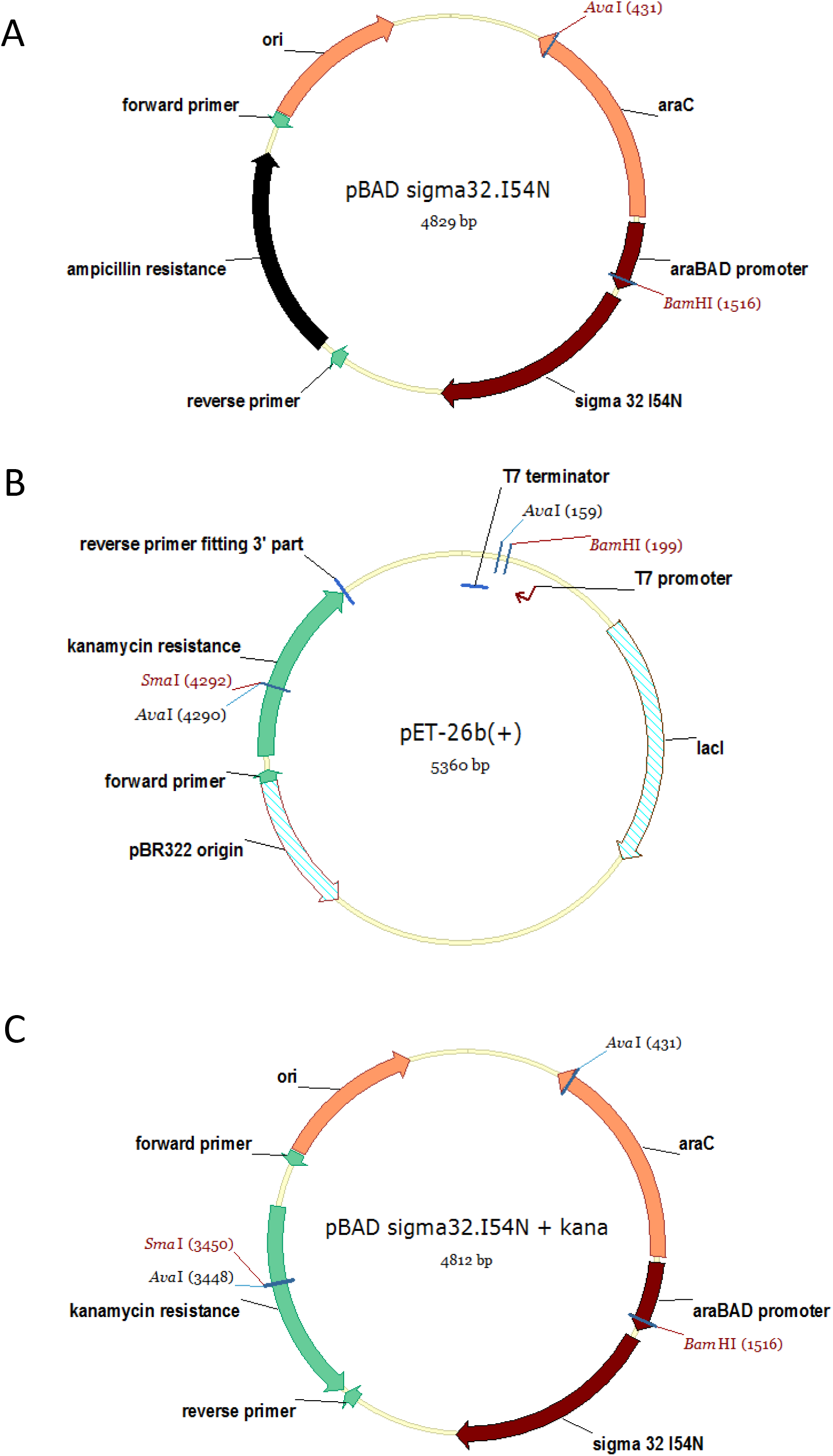
Plasmid maps. (Vector NTI, Fisher Scientific, UK). A) pBAD.Sigma32.I54N with ampicillin ARC, B) pET26b donor of the kanamycin ARC und C) the resulting pBAD.Sigma32.I54N-kana. Used restriction sites are indicated.

**Table 1.**
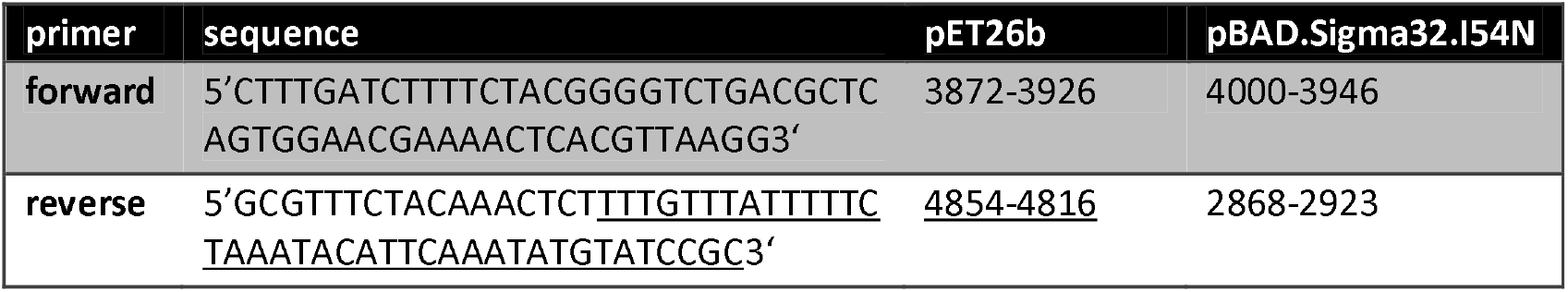
Primer for 1. PCR and their corresponding sites in vectors used.

The forward primer with 56 nucleotides matches to 100 % in both vectors and the reverse primer with 55 nucleotides fits to 71 % (39 nucleotides underlined part Table 1) to pET26b and 100 % to pBAD.Sigma32.I54N. If those identical stretches do not exist, the XL-primer can be designed by taking in 5’-3’ direction 25-30 nucleotides from the ARC flanking site of the target vector (in our case pBAD.Sigma32.I54N) combined with 25-30 nucleotides of the donor vector (in our case pET26b) to amplify first the ARC of interest and then introduce the resulting PCR product into the target vector via a second PCR. The possibility of changing the orientation of the ARC is then given via a combination of up- and downstream parts of donor and target vector, respectively, in the forward primer and vice versa for the reverse primer.

For both PCR reactions the KOD Hot Start DNA Polymerase Kit (71086, Merck, Germany) was used. The first PCR with pET26b as template and donor for the kanamycin ARC was performed as follows in a 50 μl set up: 1x KOD buffer; 2 mM MgSO_4_; 0.2 mM each of the 4 NTP’s; 0.3 μM of both primer; 0.5 ng/μl pET26b, 0.02 U/μl KOD Polymerase. The following PCR program was used:

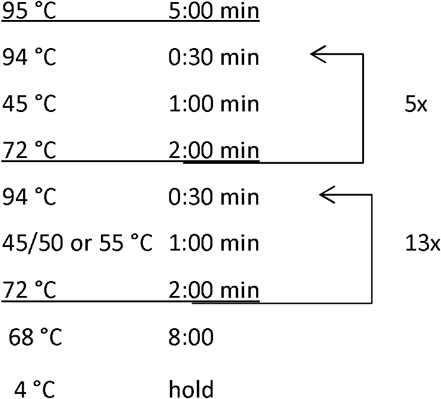

The expected PCR product of about 1000 bp was separated on a 0.8 % agarose gel (Figure 2a) and purified (LSKGEL050, Montage Gel Extraction Kit, Millipore, USA) for use as a primer pair in the final PCR step.

**Figure 2.**
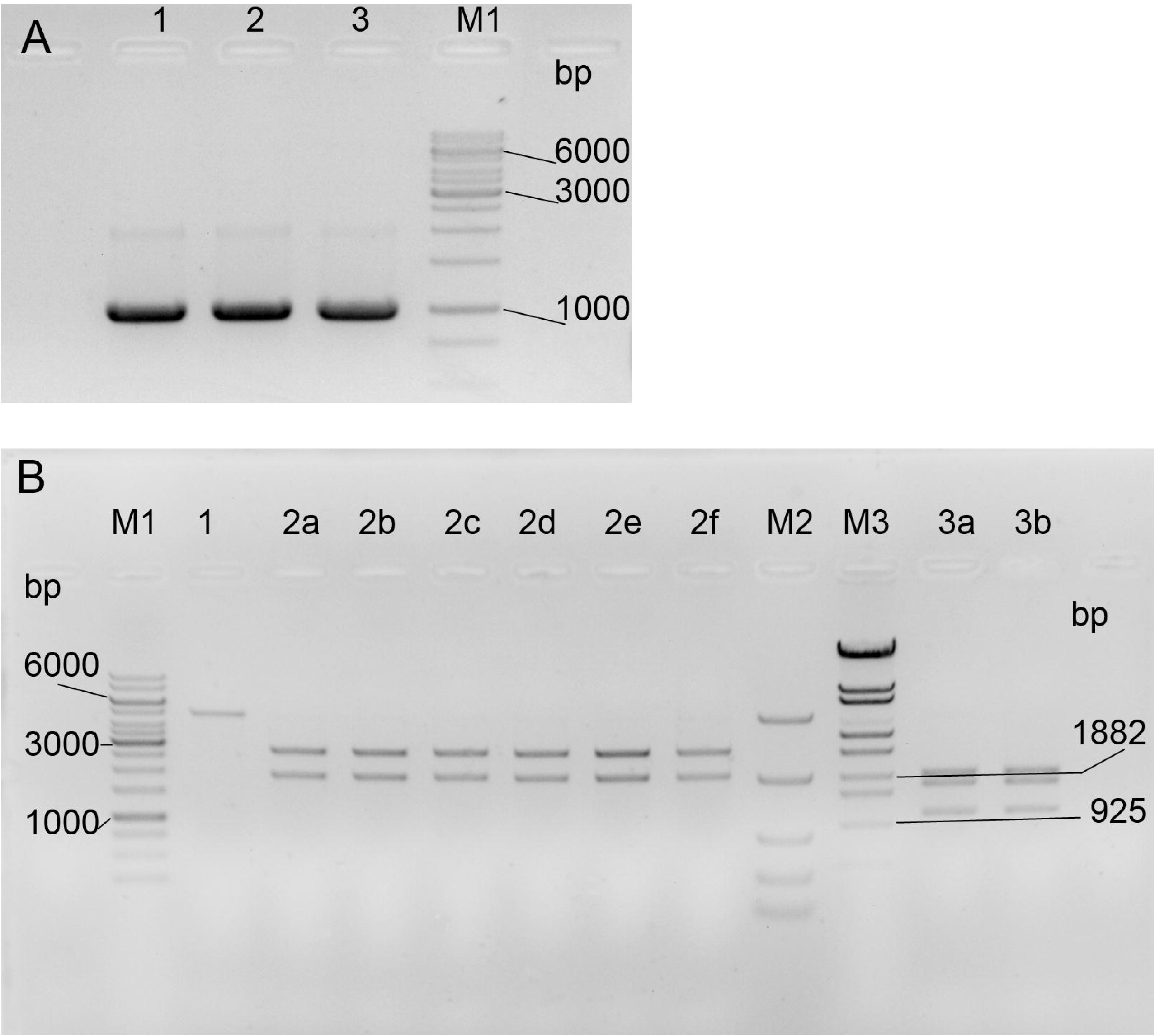
Analysis by agarose gels. A) Products of 3 parallel set ups for the first PCR with different annealing temperatures (Ta) analyzed in a 1.5 % agarose gel. Lane 1: 18 cycles with Ta 45 °C; Lane 2: 5 cycles with Ta at 45 °C, followed by 13 cycles with Ta of 50 °C; Lane 3: 5 cycles with Ta at 45 °C, followed by 13 cycles with Ta of 55 °C; M1: 1kB Marker (SM0331, Thermo Fisher Scientific) B) Restriction digest of pBAD.Sigma32.I54N and pBAD.Sigma32.I54N-kana analyzed by 0.8 % agarose gel. Marker and all “fast digest enzymes” from Thermo Fischer Scientific Lane 1: pBAD.Sigma32.I54N (4812 bp) digested with *Bam*HI (FD0054) and *Sma*I (FD0663) is only linearized as expected as it possess no cleavage site for *Sma*I; Lane 2a-f: 6 clones of pBAD.Sigma32.I54N-kana digested with BamHI and SmaI showed the expected fragments of 2878 and 1934 bp; Lane 3a und b: 2 clones of pBAD.Sigma32.I54N-kana digested with *Bam*HI and *Ava*I (FD0384, cleaves inside the kanamycin cassette and within the regulator gene araC) formed 3 fragments (1932, 1795, 1085 bp) as predicted; M1: 1kB Marker (SM0331) M2 Fast ruler Middle range with 5000, 2000, 850 400 and 100 bp (SM1113); M3 Lambda DNA/ *Eco*130I (SM0161).

The second PCR to exchange the ampicillin ARC by the amplified kanamycin ARC in pBAD.Sigma32.I54N was set up as the first PCR, but with the double amount of KOD polymerase and 6 μl of 1 kB PCR product with 40 ng/μl (final 7.2 nM) as primers. The cycling program was extended.

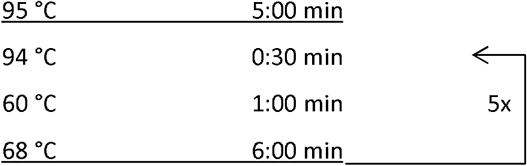

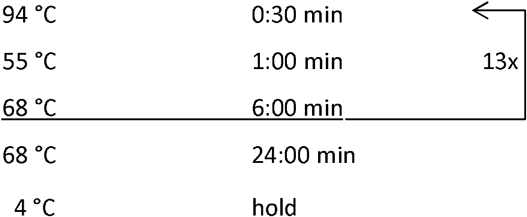

After digestion of the template plasmid with *Dpn*I at 37°C for 1 hour and heat inactivation for 5 min at 80 °C 2 μl of the treated PCR product was used to transform NovaBlue GigaSingles competent cells (71227, Merck, Germany) according to NovaBlue transformation protocol (3). Within 12 to 18 hours transformants grew on kanamycin but not on ampicillin selection plates.

In principle this result was proof enough that the kanamycin resistance cassette has been inserted correctly into the target vector. Additionally, using Sanger sequencing (Source Bioscience, UK) we demonstrated that pBAD.Sigma32.I54N-kana still carries the gene for Sigma factor.

Furthermore we analyzed plasmid preparations of six clones by restriction digest (Figure 2B).

The robustness of the method is very good, as the first PCR was successful in a broad window of annealing temperatures, many transformants were produced, six analyzed clones possessed all the expected digestion patterns and code for sigma32. An unwanted CT doubling in the middle of the reverse primer had not influenced the result negatively, as untranslated flanking regions were used as primer pairing positions and some mismatches are tolerated.

## Author contributions

N.C. performed the experiments. A.D. design the work, including the primer. N.C and AD wrote the manuscript.

## Acknowledgement

pBAD.Sigma32.I54N was a gift from Jeffery Kelly (Addgene plasmid # 59982). We thank Catherine L. Worth (FMP) for reading the manuscript.

